# Correlates of hybridization in plants

**DOI:** 10.1101/726323

**Authors:** Nora Mitchell, Lesley G. Campbell, Jeffrey R. Ahern, Kellen C. Paine, Aelton B. Giroldo, Kenneth D. Whitney

## Abstract

Hybridization is a biological phenomenon increasingly recognized as an important evolutionary process in both plants and animals, as it is linked to speciation, radiation, extinction, range expansion and invasion, and allows for increased trait diversity in agricultural and horticultural systems. Estimates of hybridization frequency vary across taxonomic groups, and previous work has demonstrated that some plant groups hybridize more frequently than others. Here, we ask on a global scale whether hybridization is linked to any of 11 traits related to plant life history, reproduction, genetic predisposition, and environment or opportunity. Given that hybridization is not evenly distributed across the plant tree of life, we use phylogenetic generalized least squares regression models and phylogenetic path analysis to detect statistical associations between hybridization and plant traits at both the family and genus levels. We find that perenniality and woodiness are each associated with an increased frequency of hybridization in univariate analyses, but path analysis suggests that the direct linkage is between perenniality and increased hybridization (with woodiness having only an indirect relationship with hybridization via perenniality). Associations between higher rates of hybridization and higher outcrossing rates, abiotic pollination syndromes, vegetative reproductive modes, larger genomes, and less variable genome sizes are detectable in some cases but not others. We argue that correlational evidence at the global scale, such as that presented here, provides a robust framework for forming hypotheses to examine and test drivers of hybridization at a more mechanistic level.

**IMPACT SUMMARY:** Although historically thought of as rare, inter-specific mating is increasingly recognized as an important evolutionary process. Hybridization can generate increased genetic and morphological variation and has been tied to increased diversification and other biological phenomena such as geographic range expansion and the success of invasive species. Here, we examine hybridization of plants on a global scale. Previous work has demonstrated that some plant groups hybridize more than others, but the reasons for this pattern remain unclear. We combine data from eight regional floras with trait data to test for associations between hybridization and different aspects of plant biology, such as life history, growth form, reproduction, and opportunity, all while accounting for the fact that plant lineages are related to each other.

We find that plant groups that are dominated by perennial species and species with woody growth forms tend to hybridize more than those dominated by annual or herbaceous species. We also find some evidence that frequent hybridization is found in plant families that are predominantly pollinated abiotically (such as by wind or water) or have higher rates of outcrossing, plant genera that have less variable genome sizes, and plant groups (both genera and families) that can reproduce asexually and have larger genome sizes. This study provides the first analysis of the global correlates of hybridization in plants. Although this correlational evidence does not provide any mechanistic explanations for these patterns, the trends we find are novel in terms of both geographic and taxonomic sale. The correlations detected provide robust hypotheses for understanding the conditions for hybridization and its contributions to evolution.

## INTRODUCTION

Hybridization is increasingly recognized as an important evolutionary phenomenon in plants (Mallet 2005; Arnold and Arnold 2006; Whitney *et al*. 2010), animals (Mallet 2005; Schwenk *et al*. 2008), and fungi (reviewed in (Albertin and Marullo 2012). Hybridization has been linked to important processes such as evolution and diversification (Anderson and Stebbins 1954; Seehausen 2004), adaptive radiation (Anderson and Stebbins 1954; Stebbins 1959; Barton 2001; Seehausen 2004; Yakimowski and Rieseberg 2014, Marques *et al*. 2019), and speciation (Rieseberg 2003; Mallet 2007; Rieseberg *et al*. 2007; Soltis and Soltis 2009; Abbott *et al*. 2013). Hybridization has enabled plant breeders to transfer desirable traits among species for both agricultural and horticultural purposes (Allard 1999). In contrast, hybridization has also been linked to numerous conservation concerns such as biological invasion (Ellstrand and Schierenbeck 2000; Schierenbeck and Ellstrand 2009; Whitney *et al*. 2010; Hovick *et al*. 2012; Hovick and Whitney 2014), escape of novel traits via crop-wild hybridization (Ellstrand and Hoffman 1990; Zapiola *et al*. 2008), and even extinction via hybridization (Rhymer and Simberloff 1996; Wolf *et al*. 2010; Todesco *et al*. 2016, Campbell *et al*. In Press). A deep understanding of hybridization is thus necessary to understand evolutionary principles, to provide for agricultural needs, and to inform conservation management decisions.

There is evidence for hybridization in unexpected situations, for instance between distantly related species (Rothfels *et al*. 2015) or in cases of cryptic hybridization with molecular but little morphological evidence (Cronn and Wendel 2004; Soltis *et al*. 2007; McIntosh *et al*. 2014; Mitchell and Holsinger 2018). Focke (1881, *in* Stebbins 1959 and Grant 1981) first made the observation that rates of hybridization differ across plant taxa. More modern analyses based on floras or surveys of the literature have found different rates of hybridization in different taxonomic groups, with evidence for phylogenetic signal (Ellstrand *et al*. 1996; Whitney *et al*. 2010; Abbott 2017; Beddows and Rose 2018). Ferns and their allies and specific flowering plant families (such as Orchidaceae, Lamiaceae, Asparagaceae, and Asteraceae) contain high numbers of hybridizing species, while other families appear to contain few hybrids (such as Caryophyllaceae, Cyperaceae, and Apiaceae) (Whitney *et al*. 2010).

Hypotheses as to why some groups hybridize more than others center on traits related to life history, reproduction, genetics, and opportunity or environment. Researchers have either advanced theoretical reasons for a connection between a trait and increased hybridization, or have identified correlational evidence to support a connection without a theoretical justification (summarized in Table 1 and expanded on in Table S1). These traits may be associated with the *formation* of hybrids, i.e. allowing for interspecific mating and production of offspring, or may be associated with the *persistence* of hybrids, i.e. allowing for the continued propagation of a hybrid lineage after formation. Briefly, we expected that plant groups dominated by perennial species (Grant 1958, 1981; Stace 1975; Ellstrand *et al*. 1996; Beddows and Rose 2018) or woody species (Stebbins 1959; Beddows and Rose 2018) will contain more hybrids than those dominated by annual or herbaceous life histories, because longer lifespans associated with perenniality and woodiness may allow hybrid individuals to produce offspring over time despite partial sterility, allowing for persistence of these hybrid lineages (Ellstrand *et al*. 1996). We also expected higher rates of hybridization in plant groups with traits that increase the likelihood of interspecific mating, either by reducing barriers to gene flow or promoting outbreeding. These include traits such as pollination syndrome (contrasting evidence for increased hybridization with both biotic: Rieseberg and Wendel 1993, or abiotic: Ellstrand et al. 1996, pollination syndromes), bilaterally symmetrical flowers (Stebbins 1959, Sargent 2004), reproductive systems that require cross-breeding (higher outcrossing rates: Stace 1975; Grant 1981), sexual breeding systems (Grant 1981), and generative/non-vegetative reproductive systems (Ellstrand *et al*. 1996). Some groups may be genetically predisposed to hybridize, for instance lineages with few chromosomal translocations which allow for greater fertility in hybrids (Grant 1981), smaller genome sizes (as reported in Bureš *et al*. 2004), or less variable genome sizes which may allow for greater interspecific compatibility. Finally, hybridization may be the product of opportunity, where greater opportunity might be conferred via having agricultural relatives that by nature are abundant and widespread, being less threatened by extinction, or being found in more disturbed environments where contact with relatives might be initiated (Anderson and Stebbins 1954; Grant 1981; Guo 2014).

**Table 1.**
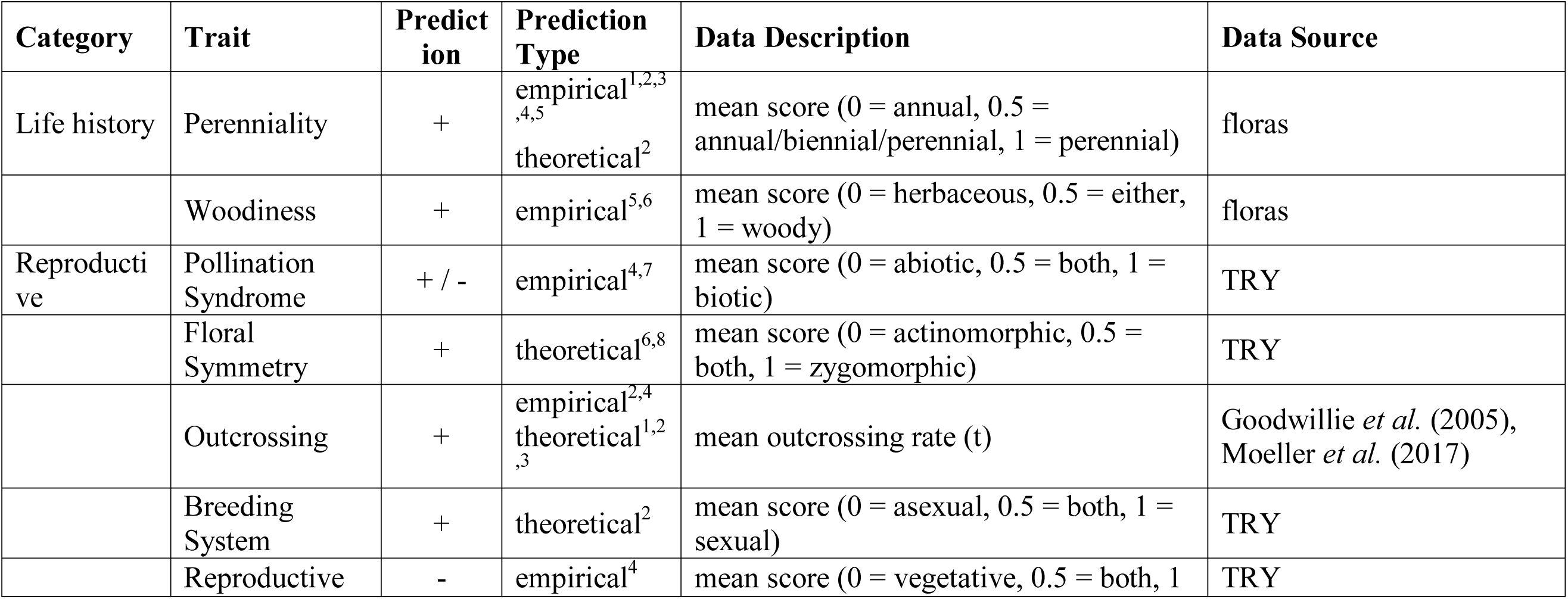
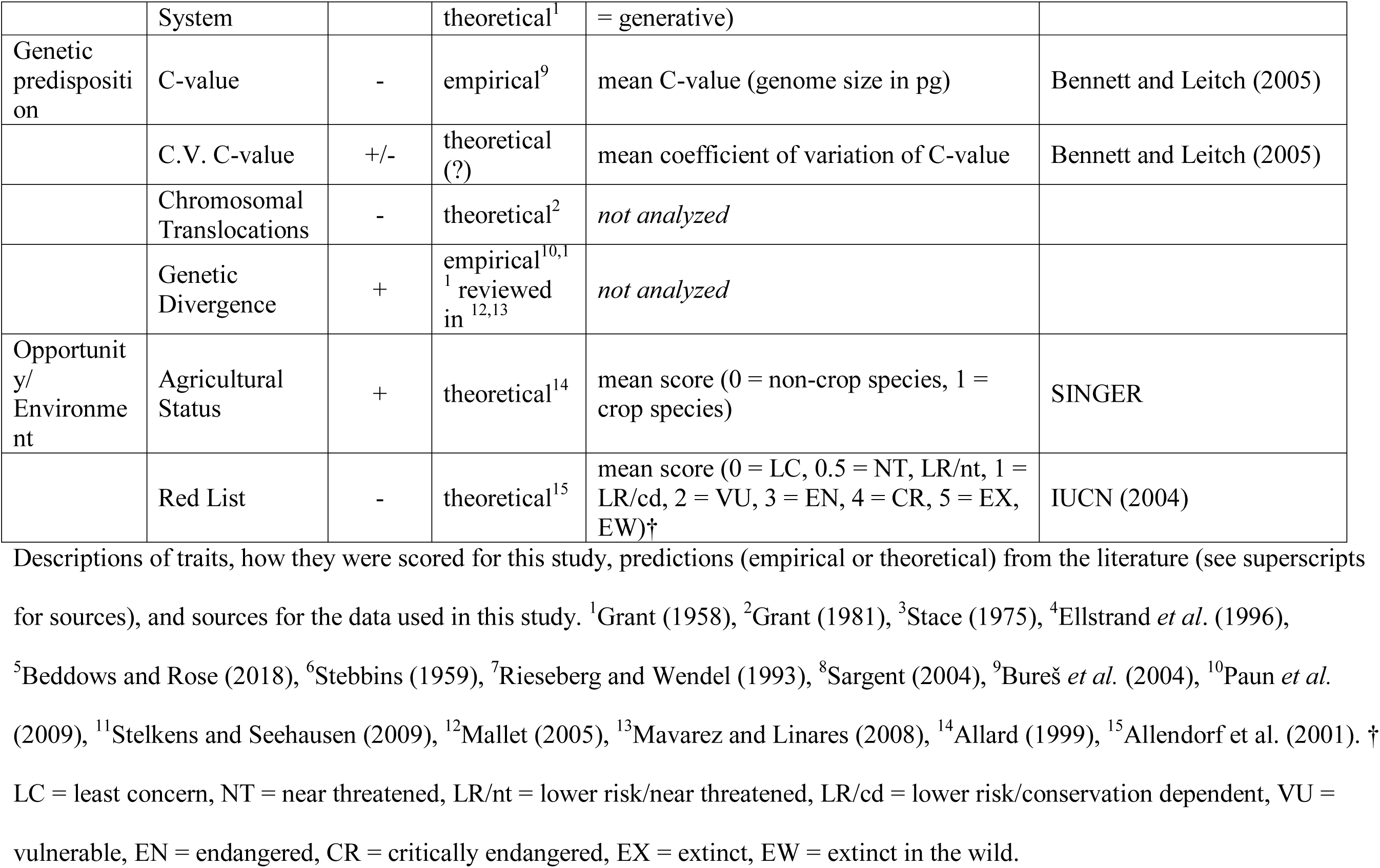
A review of the potential traits associated with hybridization in plants, as identified by a literature search, with further information on data types and sources in our analysis. The “Prediction” column gives the predicted sign of the association between the trait and hybridization propensity, relative to the orientation in the “Data Description” column. “Prediction Type” distinguishes whether predictions from the literature are based on a theoretical argument or simply on an observed (but not phylogenetically corrected) empirical association. We expand on proposed mechanisms in Table S1. Data used in analyses were mean scores across all species within the group of interest (family or genus). When we did not have data to test the potential relationship, the “Data Source” column is blank.

At the regional scale, measures of hybridization have been empirically linked to various plant attributes. Beddows and Rose (2018) performed a case study on the flora of Michigan, a single state in the United States. They surveyed the published flora for interspecific hybrids and several plant attributes, including life history and life form, and used multiple logistic regressions to determine what factors were correlated with various measures of hybridization. Although taxonomic order was included in the analysis, they did not explicitly account for the phylogenetic non-independence of the taxa analyzed. In their analysis, hybridization was positively correlated with perenniality, woodiness, habitat disturbance, and number of herbarium records, and they additionally detected significant effects of taxonomic order (Beddows and Rose 2018).

Thus far, there has been no comprehensive analysis of the potential correlates of hybridization in plants at the global scale, nor has there been an analysis accounting for phylogenetic non-independence among taxa. Here, we build on the work of Whitney *et al*. (2010), which quantified hybridization across the globe in 282 different plant families and 3212 genera using data from eight regional floras. We expanded this dataset and combined it with trait data collected from the regional floras and additional external datasets to ask whether hybridization in plants (quantified using two metrics) is statistically associated with 11 different traits at both the family and genus levels, while simultaneously accounting for the phylogenetic non-independence of the taxa analyzed.

## METHODS

### Extent of hybridization

To characterize the extent of hybridization across vascular plant families, we analyzed eight floras: the Great Plains of the U.S. (McGregor and Barkley 1986), the British Isles (Stace 1997); Hawai’i (Wagner *et al*. 1999); the Intermountain Region of the western U.S. (Cronquist *et al*. 1972); the Northeastern U.S. (Magee and Ahles 1999); California (Hickman 1993); Europe (Tutin *et al*. 1964); and Victoria, Australia (Walsh and Entwisle 1994) (Fig. 1). These floras are the same as those used in Whitney *et al*. (2010), with the exception that we have here included the final published volume of the Intermountain Region (volume 2A, 2012). Floras were chosen nonrandomly to include those that contained multiple mentions of hybrids, and are therefore a biased subset reflecting regions where hybrids are common or, more likely, reflecting authors interested in hybridization and attuned to recording instances of it.

**Fig. 1.**
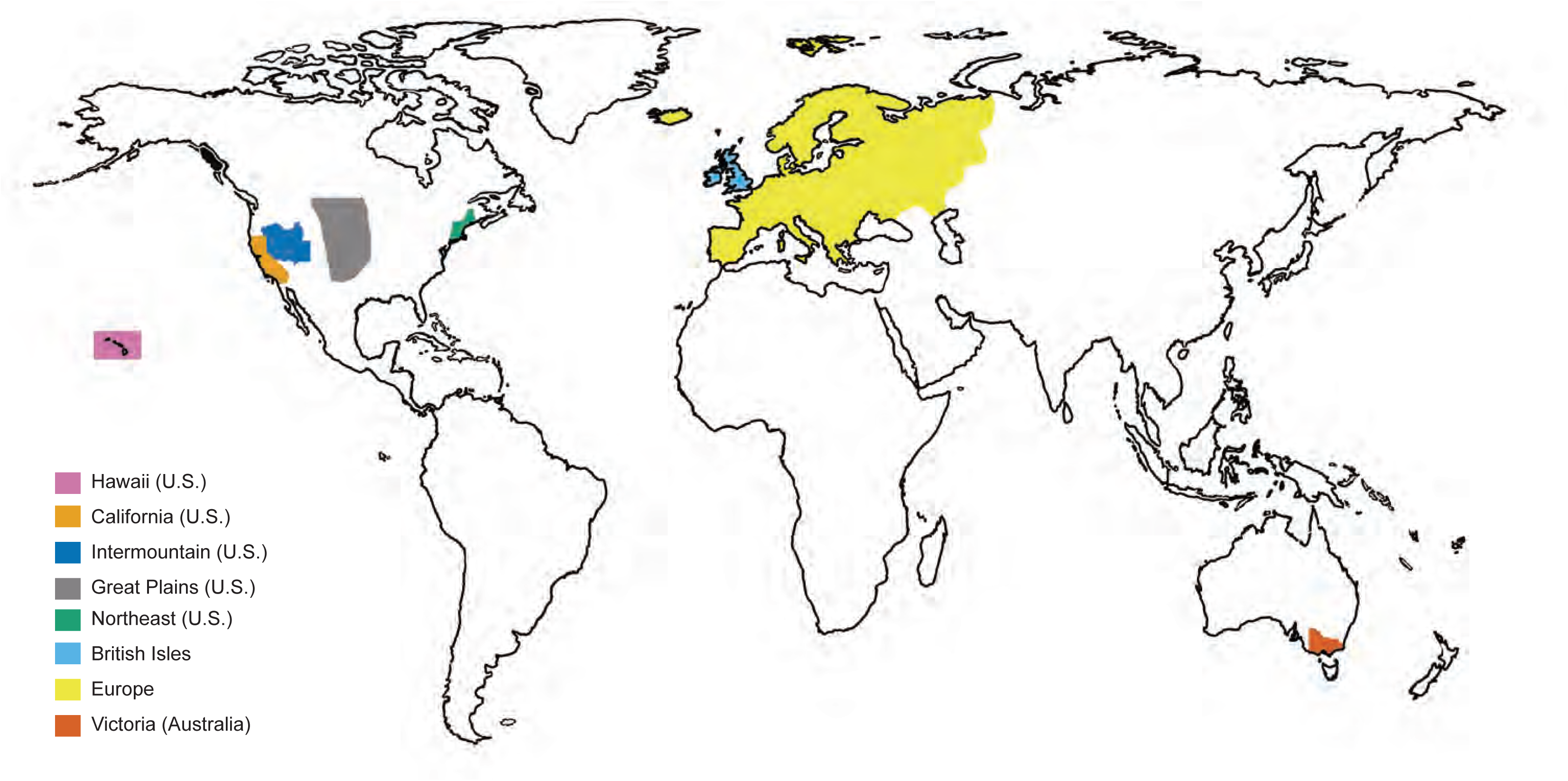
World map indicating the coverage areas of the floras used in this study. Pink = Hawaii (U.S.), light orange = California (U.S.), dark blue = Intermountain (U.S.), gray = Great Plains (U.S.), green = Northeast (U.S.), light blue = British Isles, yellow = Europe, dark orange = Victoria (Australia).

For each vascular plant family in each flora, the numbers of interspecific hybrids and the numbers of non-hybrid species were determined as in Whitney *et al*. (2010). For counting purposes, we follow Ellstrand *et al*. (1996) in defining a “hybrid” as a hybrid type derived from a unique combination of two parental species. Thus, in each flora, each pair of hybridizing species was counted as generating a single hybrid, even if there was evidence that the pair had hybridized multiple times. Our recognition of an interspecific hybrid does not imply that it was formally or taxonomically recognized in the flora (though some were), nor does it imply processes such as hybrid speciation or the formation of a hybrid population that is stable over the long-term. It simply is an observation that a pair of parental species has interbred and resulted in hybrid offspring that have persisted in the wild long enough to be noted by an author of a flora. Only native and naturalized taxa were considered. Taxa clearly resulting from anthropogenic crosses (e.g. “garden hybrids”) and taxa only in cultivation were ignored. We tallied intra- and inter-generic hybrids separately, and the latter were split between genera (e.g., half of each hybrid was assigned to each contributing genus). We did not count hybrids among subspecies or probable primary intergradation (diverging sub-populations maintaining genetic connections, Stebbins 1959). In each flora, each pair of hybridizing species was counted as generating a single hybrid taxon, even if there was evidence that the pair had hybridized multiple times. We also counted naturalized hybrids mentioned in a flora that apparently arose outside the region covered by the flora. Finally, in some floras, particular groups were described as producing multiple hybrids without detailed specification of their numbers or the parental species involved. In these few cases we estimated the number of hybrids as either 2 hybrids or 20% of the number of species present, whichever was greater. We analyzed all floras at the generic level and reassigned those genera (with their associated counts of species and hybrids) to families based on The Plant List (http://www.theplantlist.org/) to accommodate taxonomic changes since the publication of the floras.

We collected hybridization data on 282 plant families and 3229 different genera. Observations of genera with a single non-hybrid species identified in a single flora were then eliminated to avoid including groups with no chance for hybridization, and a single family that could not be placed phylogenetically with confidence (Capparaceae, see below) was also excluded. This resulted in a final sample size of 195 families for the family-level analysis. For the genus-level analysis, we were unable to place 34 genera in the phylogeny (see below), resulting in a final sample size of 1772 genera (Table S2).

We characterized hybridization for each family or group using two metrics: hybridization propensity and hybrid ratio, for completeness and comparability. Hybridization propensity reflects the realized percentage of all possible hybrid combinations and is calculated as in Whitney *et al*. (2010). For a taxonomic group of *n* nonhybrid species:

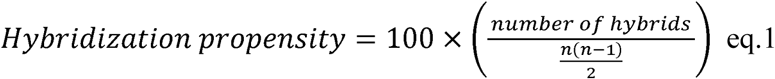

Although it is unrealistic that every pair of species within a group hybridizes (so the denominator of eq. 1 is perhaps unrealistically large), we feel that bounds on the percentage of species that could potentially hybridize would require additional information beyond the scope of this study. Hybrid ratio, employed by Beddows and Rose (2018), reflects the number of hybrid combinations relative to all nonhybrid taxa. For a taxonomic group of *n* nonhybrid species:

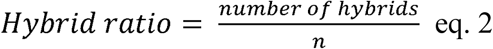

We calculated and analyzed both to be able to compare our findings to previous studies. Note the scale difference: by convention, hybridization propensity is a percentage bounded between 0 and 100, while hybrid ratio is unbounded (in practice, it ranges from 0 – 0.15 with outliers up to 1.2). For each genus, numbers of both nonhybrids and hybrids were calculated by summing hybrid counts across all floras analyzed. No attempt was made to avoid ‘double counting’ of hybrids formed from the same parents in different regions. Thus, each metric incorporates information on both the number of hybridizing taxa and the frequency with which they hybridize in different regions. Genus-level metrics were calculated based on the observations across all floras, while family-level metrics were weighted means of metrics of the component genera (weighted by species number in each genus). Both hybridization propensity and hybrid ratio measures were log-transformed prior to analysis to more closely match assumptions of normality.

### Traits of plant groups

The number of annual, biennial, and perennial species, and the number of herbaceous vs. woody species, were summed for each genus in each flora. The floras provided remarkably complete data on these variables (>95% species covered), but missing data on perenniality and woodiness of the species were determined from other sources (e.g. USDA plants database). Species described as intermediate (e.g. “annual/biennial”) were split between categories (e.g. counted as 0.5 annuals and 0.5 biennials). Species were considered woody if they were characterized by substantial aboveground woody biomass, e.g. “trees”, “shrubs”, “subshrubs”, “woody vines” and “lianas”. Species with rootstocks as the only woody parts were considered herbaceous. For genus and family-level analyses, we used the percentage of species scored as perennial and the percentage of species scored as perennial as our trait data (Table 1).

Data for several traits were downloaded from the TRY database (Kattge *et al*. 2011). These included information on pollination syndrome (abiotic or biotic: Giroldo 2016; Fitter and Peat 1994; Koike 2001; Ogaya and Peñuelas 2003; Diaz *et al*. 2004; Kühn *et al*. 2004; Gachet *et al*. 2005; Moretti and Legg 2009; Onstein *et al*. 2014; de Frutos *et al*. 2015; Chapin unpubl.; Leishman unpubl.), breeding system (asexual or sexual: Kühn *et al*. 2004), floral symmetry (actinomorphic or zygomorphic: Dressler *et al*. 2014), and reproductive system (vegetative or generative: Fitter and Peat 1994; Kühn *et al*. 2004; Klimešová and de Bello 2009). For each species in the TRY dataset, trait values were simplified to be either 0, 0.5 (for mixed or combined), or 1 (see Table 1 for coding schemes for individual traits). We used genus or family-level means for pollination syndrome, breeding system, floral symmetry, and reproductive system as trait data in subsequent analyses.

We compiled additional trait data from other sources. We assessed agricultural status by calculating the percentage of species in each family that were listed as crop species as defined in the System-wide Information Network for Genetic Resources database (http://singer.cgiar.org/Search/SINGER/search.htm, downloaded July 2009). We assessed threatened status using data from the Red List (Baillie *et al*. 2004). We assigned numeric values representing each species’ threatened status (see Table 1 for scoring categories) and used genus- or family-level means. We estimated genus- and family-level mean outcrossing rates from Goodwillie *et al*. (2005) and Moeller *et al*. (2017). Finally, genome size estimates (both “Prime Estimates” and others) were downloaded from the Plant DNA C-values database (Bennett and Leitch 2005). We calculated the mean genome size per species (including all ploidy level variants, if present in the database) and then calculated genus and family-level means. C-value was log-transformed prior to analysis. We also estimated the coefficient of variation for genome size by calculating mean c-values for each ploidy level of each species, then calculating the coefficient of variation across these means for each genus and family levels. See Table 1 for full information on the traits assessed.

### Composite tree construction and phylogenetic signal

Subsequent analyses were conducted in R v3.3.3 (R Core Development Team 2016). To account for the phylogenetic nonindependence of our observations, we used phylogenetic generalized least squares regression (PGLS regression: Grafen 1989; Martins and Hansen 1997). The family-level seed plant phylogeny was imported from the tree of Qian and Jin (2016) (an updated and corrected version of Zanne *et al*. 2014) into R using the “ape” package (Paradis *et al*., 2004). The phylogeny was trimmed and resolved to include only the seed plant families for which we had data using the S.Phylomaker function from Qian and Jin (2016). To include non-seed plants, we manually constructed phylogenies in Mesquite v3.40 (Maddison and Maddison 2018) based on their position in the literature for ferns (Smith *et al*. 2006) and fern allies (Pryer *et al*. 2004) and combined them in R. To construct a genus-level phylogeny, we used S.Phylomaker and added within-family relationships for the ferns and their allies by hand based on the literature (Hauk *et al*. 2003; Pryer *et al*. 2004; Schneider *et al*. 2004a,b; Ebihara *et al*. 2006; Liu *et al*. 2007; He and Zhang 2012; Sundue *et al*. 2014; de Gasper *et al*. 2017). Phylogenies are available from the lead author on request.

We estimated phylogenetic signal via Pagel’s λ separately for each measure of hybridization and each trait using the *phylopars()* function with model set to “lambda” in the “Rphylopars” package (Goolsby *et al*. 2017). We compared this model to a star phylogeny with lambda = 0 using likelihood ratio tests. Although the “Rphylopars” package allows imputation of missing trait values (Bruggeman *et al*. 2009; Goolsby *et al*. 2017), we had high amounts of missing data (for a given trait, up to 61% in families and 89% in genera) so chose instead to prune trees to exclude taxa with missing data before each analysis.

### Analyses of hybridization vs. potential correlates

We calculated the raw correlations between all 11 traits and the two hybridization metrics at both the family and generic levels using the *corr.test()* function in the R package “psych” (Revelle 2017). However, raw correlations do not account for phylogenetic non-independence among taxa (Felsenstein 1985) so we report these only for frame of reference.

Phylogenetic generalized least squares (PGLS) regression provides a flexible framework for detecting associations among traits under different evolutionary models (Grafen 1989; Martins and Hansen 1997). PGLS was conducted using the *phylopars.lm()* function in the R package “Rphylopars” (Goolsby *et al*. 2017). We performed univariate PGLS regressions for each of our traits on both metrics of hybridization at the family and generic levels, subsetting the data and phylogenies to prevent imputation (see above for explanation). Note that we were missing values for some traits due to lack of available data and for other traits because they were not applicable to all taxonomic groups (e.g., only seed plants have pollination syndromes, and only flowering plants have florals symmetry). Regressions were performed under the Brownian Motion (BM), Ornstein-Uhlenbeck (OU), and early burst (EB) models of evolution, and then compared using AIC and BIC. As either BM or EB was the best model across all traits, and as all models were within 2 AIC, we report BM results as representative. We corrected for multiple comparisons using the Benjamini-Hochberg procedure (Benjamini and Hochberg, 1995) within each hybridization measure and taxonomic level combination (11 total tests per combination), using a false discovery rate of 0.05.

### Phylogenetic path analysis

A potential multivariate analysis including all 11 traits as predictors of hybridization was not practical, because of missing trait data. However, we did have nearly complete information for woodiness and perenniality. In order to simultaneously estimate the relationships between hybridization and both perenniality and woodiness, we used the “phylopath” package (van der Bijl 2018) to run phylogenetic path analyses. Although causal relationships cannot be determined from correlational evidence, path analysis allows for an understanding of direct and indirect relationships under proposed causal models (von Hardenberg and Gonzalez-Voyer 2013; Kennedy *et al*. 2018). We used these models to determine the relative strength of these two highly correlated predictors of hybridization when present in the same model. We tested five path structures for each combination of taxonomic level and measure of hybridization (Fig. S1). The fit of models was estimated using the C statistic, which provides an estimate of goodness of fit of the model to the data (Shipley 2013). We report results from the best model using CICc, the C statistic information criterion (von Hardenberg and Gonzalez-Voyer 2013).

## RESULTS

### Hybridization metrics and phylogenetic signal

In the 195 plant families analyzed, 112 contained hybrids and 83 did not. The mean value for family-level hybridization propensity was 2.55% (range = 0 – 100%) and for hybrid ratio was 0.086 (range = 0 – 1.196) (Fig. 2, Table S3). At the family level, the log-transformed values for hybridization propensity and hybrid ratio were significantly correlated (corr = 0.701, p < 0.001) (Table S4). There was significant phylogenetic signal in hybridization propensity (λ = 0.30, p < 0.001) and a lower, but still significant, measure of phylogenetic signal in hybrid ratio (λ = 0.14, p < 0.01) (Table 2). Eight out of 11 traits had significant phylogenetic signal at the family level (perenniality, woodiness, percent agricultural, floral symmetry, pollination syndrome, reproductive system, C-value, and coefficient of variation in C-value; see Table 2).

**Fig. 2.**
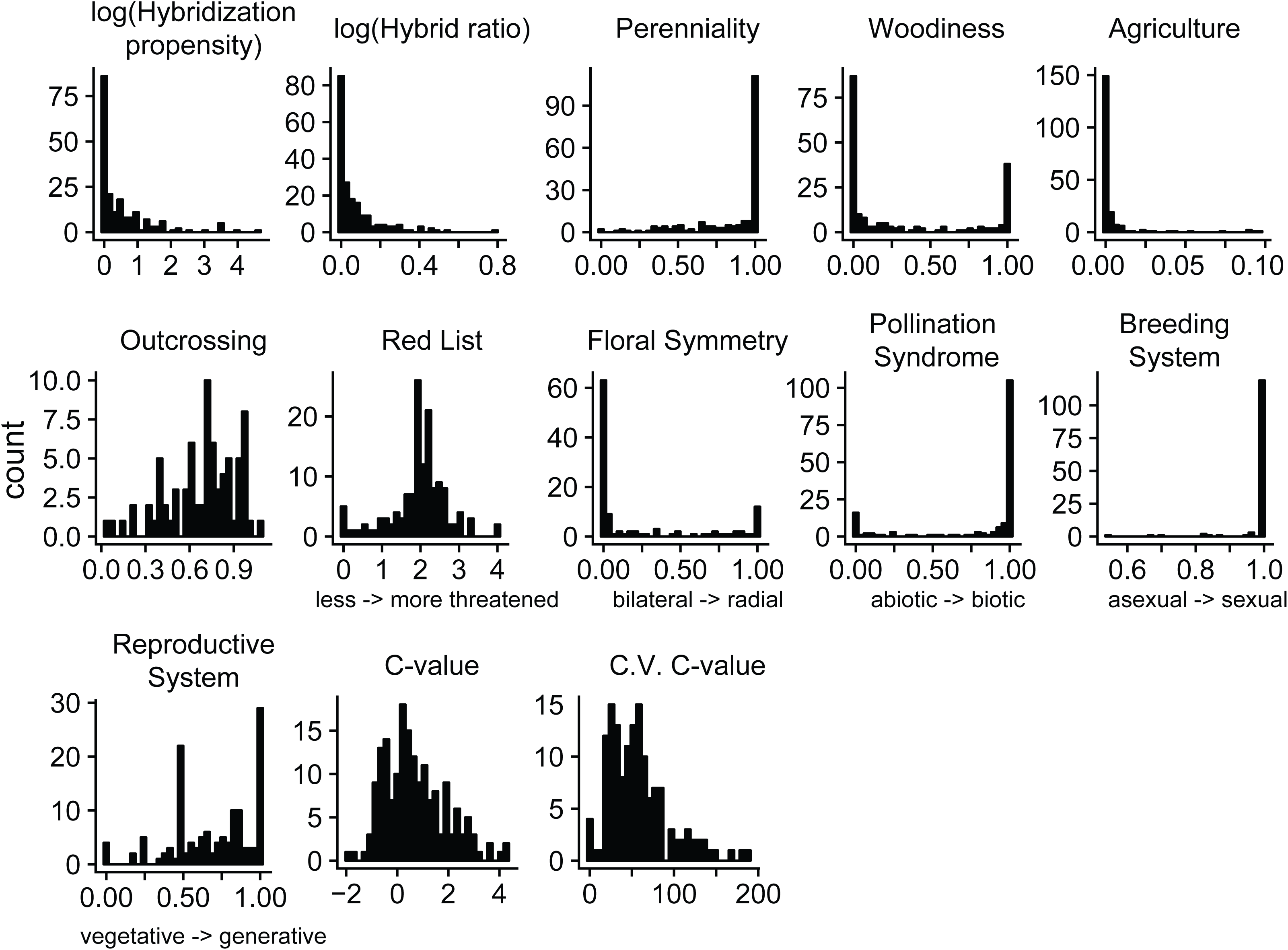
Distributions of family-level hybridization metrics and family-average traits. See Table 2 for trait descriptions and units. Non-intuitive trait values have brief descriptions on the x-axes.

**Table 2.** Phylogenetic signal (Pagel’s λ and associated chi-square statistics and p-values) of hybridization measures and potential predictors at different taxonomic levels.

|  | Family Level |  |  |  |  |  | Genus Level |  |  |  |  |
| --- | --- | --- | --- | --- | --- | --- | --- | --- | --- | --- | --- |
| Trait | N observed | Pagel's $\lambda$ | Chi-Square | DF | P-value | | N observed | Pagel's $\lambda$ | Chi-Square | DF | P-value |
| Hybridization Propensity | 195 | 0.30 | 32.31 | 1 | 0.000 |  | 1772 | 0.11 | 52.28 | 1 | 0.000 |
| Hybrid Ratio | 195 | 0.14 | 8.06 | 1 | 0.005 |  | 1772 | 0.13 | 52.05 | 1 | 0.000 |
| Perenniality | 195 | 0.22 | 10.34 | 1 | 0.001 |  | 1754 | 0.47 | 314.01 | 1 | 0.000 |
| Woodiness | 195 | 0.47 | 40.87 | 1 | 0.000 |  | 1767 | 0.80 | 968.73 | 1 | 0.000 |
| Percent Agricultural | 195 | 0.26 | 3.90 | 1 | 0.048 |  | 1772 | 1.00 | 6738.41 | 1 | 0.000 |
| Outcrossing | 76 | 0.01 | 0.01 | 1 | 0.943 |  | 158 | 0.24 | 3.72 | 1 | 0.054 |
| Red List | 138 | 0.00 | -0.01 | 1 | 1.000 |  | 374 | 0.25 | 21.45 | 1 | 0.000 |
| Floral Symmetry | 114 | 0.51 | 13.33 | 1 | 0.000 |  | 235 | 0.76 | 124.51 | 1 | 0.000 |
| Pollination Syndrome | 164 | 0.79 | 70.89 | 1 | 0.000 |  | 878 | 0.93 | 1208.71 | 1 | 0.000 |
| Breeding System | 130 | 0.03 | 0.17 | 1 | 0.678 |  | 639 | 0.09 | 8.87 | 1 | 0.003 |
| Reproductive System | 133 | 0.32 | 18.48 | 1 | 0.000 |  | 655 | 0.46 | 135.09 | 1 | 0.000 |
| C-value | 177 | 0.66 | 57.11 | 1 | 0.000 |  | 761 | 0.74 | 476.77 | 1 | 0.000 |
| C.V. C-value | 144 | 0.37 | 7.04 | 1 | 0.008 |  | 522 | 0.00 | -0.00 | 1 | 1.000 |

We analyzed 1772 different plant genera, of which 492 contained hybrids and 1280 did not. The mean value for genus-level hybridization propensity was 2.885% (range = 0 – 300%) and for hybrid ratio was 0.060 (range = 0 – 1.609) (Table S3). At the genus level, the log-transformed values for hybridization propensity and hybrid ratio were significantly correlated (corr = 0.846, p < 0.001) (Table S4). We also detected low but significant phylogenetic signal in hybridization propensity (λ = 0.11, p < 0.001) and hybrid ratio (λ = 0.13, p < 0.001) at the genus level (Table 2). Nine out of 11 traits had significant phylogenetic signal at the genus level (all but outcrossing and the coefficient of variation of C-value; see Table 2).

### Plant traits

We assessed 11 potential correlates of hybridization using data from the floras as well as other sources (Table 1). The dataset was dominated by perennial and herbaceous taxa as well as by taxa with radially symmetric flowers, biotic pollination syndromes, sexual breeding systems, and generative reproductive systems (Fig. 2, Table S3).

### Correlates of hybridization

Using univariate regressions at the family level, we detected significant associations (p < 0.05) linking abiotic pollination syndrome to increased hybridization propensity and a trend (0.05 < p < 0.10) for links between both higher outcrossing rates and larger genome sizes and hybridization propensity (Fig. 3, Table 3). We detected associations between perenniality, woodiness, and more abiotic pollination syndromes with hybrid ratio, although only the latter was significant (Fig. 3, Table 3). However, after correcting for multiple comparisons, none of these family level associations were significant. Adjusted R^2^ values were very low, with a maximum of 0.034.

**Fig. 3.**
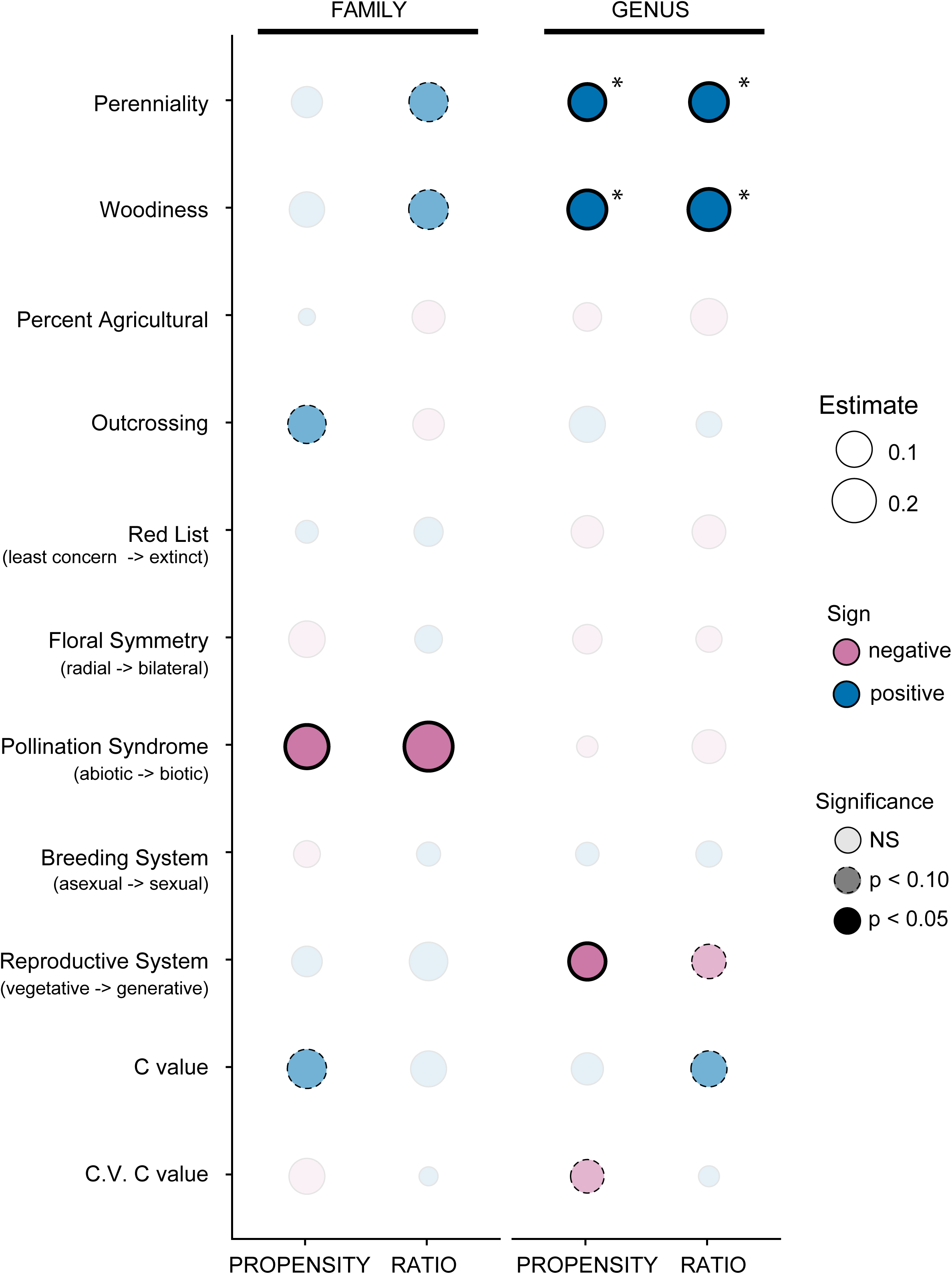
Predictors of hybridization propensity and hybrid ratio at the family (left) and genus (right) levels from PGLS univariate regressions. Sizes of the circles indicate the absolute value of the strength of the estimate. Color indicates the sign (positive = blue, negative = pink) of the estimate. The transparency and border indicate the significance of the estimate: lightest shaded circles were not significant (p > 0.10), medium shading with dashed borders indicates a trend (p < 0.10), and darkest shading with solid bold borders indicates statistical significance (p < 0.05). Asterisks indicate that the relationship is significant after a Benjamini-Hochberg procedure.

**Table 3.** Univariate PGLS results at different taxonomic levels. * indicates relationships significant after Benjamini-Hochberg procedure, raw p-values < 0.05 are in bold.

|  | Family Level |  |  |  |  |  |
| --- | --- | --- | --- | --- | --- | --- |
|  | Hybridization Propensity |  |  | Hybrid Ratio |  |  |
| Trait | Estimate | P-value | Adjusted-R<sup>2</sup> | Estimate | P-value | Adjusted-R<sup>2</sup> |
| Perenniality | 0.057 | 0.391 | -0.001 | 0.135 | 0.061 | 0.013 |
| Woodiness | 0.093 | 0.206 | 0.003 | 0.141 | 0.077 | 0.011 |
| Percent Agricultural | 0.004 | 0.951 | -0.005 | -0.072 | 0.322 | 0.000 |
| Outcrossing | 0.125 | 0.083 | 0.027 | -0.060 | 0.498 | -0.007 |
| Red List | 0.013 | 0.857 | -0.007 | 0.043 | 0.606 | -0.005 |
| Floral Symmetry | -0.105 | 0.167 | 0.008 | 0.034 | 0.744 | -0.008 |
| Pollination Syndrome | -0.191 | **0.019** | 0.028 | -0.267 | **0.010** | 0.034 |
| Breeding System | -0.029 | 0.610 | -0.006 | 0.017 | 0.817 | -0.007 |
| Reproductive System | 0.053 | 0.392 | -0.002 | 0.127 | 0.106 | 0.012 |
| C-value | 0.136 | 0.084 | 0.011 | 0.099 | 0.288 | 0.001 |
| C.V. C-value | -0.099 | 0.183 | -0.006 | 0.005 | 0.958 | -0.007 |
|  | Genus Level |  |  |  |  |  |
|  | Hybridization Propensity |  |  | Hybrid Ratio |  |  |
| Trait | Estimate | P-value | Adjusted-R<sup>2</sup> | Estimate | P-value | Adjusted-R<sup>2</sup> |
| Perenniality | 0.103 | **0.000\*** | 0.007 | 0.123 | **0.000\*** | 0.011 |
| Woodiness | 0.126 | **0.000\*** | 0.007 | 0.161 | **0.000\*** | 0.011 |
| Percent Agricultural | -0.039 | 0.840 | -0.001 | -0.109 | 0.568 | 0.000 |
| Outcrossing | 0.101 | 0.171 | 0.006 | 0.024 | 0.752 | -0.006 |
| Red List | -0.067 | 0.295 | 0.000 | -0.078 | 0.248 | 0.001 |
| Floral Symmetry | -0.045 | 0.515 | -0.002 | -0.026 | 0.698 | -0.004 |
| Pollination Syndrome | -0.009 | 0.904 | -0.001 | -0.079 | 0.314 | 0.000 |
| Breeding System | 0.015 | 0.671 | -0.001 | 0.026 | 0.515 | -0.001 |
| Reproductive System | -0.106 | **0.011\*** | 0.008 | -0.085 | 0.074 | 0.003 |
| C-value | 0.065 | 0.226 | 0.001 | 0.101 | 0.081 | 0.003 |
| C.V. C-value | -0.077 | 0.064 | 0.005 | 0.008 | 0.871 | -0.002 |

At the genus level, increased perenniality and woodiness were associated with increased hybridization in both metrics. These relationships were still significant after a Benjamini-Hochberg correction (Table 3). There was a slight association (0.05 < p < 0.10) between less variable genome sizes and increased hybridization propensity and a significant association (after correcting for multiple comparisons) between more vegetative reproductive systems and hybridization propensity. There were trends for genera with more vegetative reproductive systems and larger genome sizes to have higher values of hybrid ratio (Fig. 3, Table 3). Adjusted R^2^ values were also very low, with a maximum of 0.011. Family- and genus-level relationships were generally in consensus, in that there were no instances where a well-supported association at one taxonomic level was well-supported in the opposite direction at the other taxonomic level (Fig. 3, Table 3).

### Phylogenetic path analysis

To account for the high correlations among two traits with detectable associations with hybridization in the univariate regressions), we examined relationships between hybridization and both perenniality and woodiness using phylogenetic path analyses (Fig. S1). At both the family and genus levels, the best models indicate that woodiness does not have a direct link to hybridization, but instead has an indirect association via a pathway including perenniality and perenniality’s direct association with hybridization (Fig. 4, Table S5). The estimated path coefficients were all positive and above zero +/− standard error (Table S6).

**Fig. 4.**
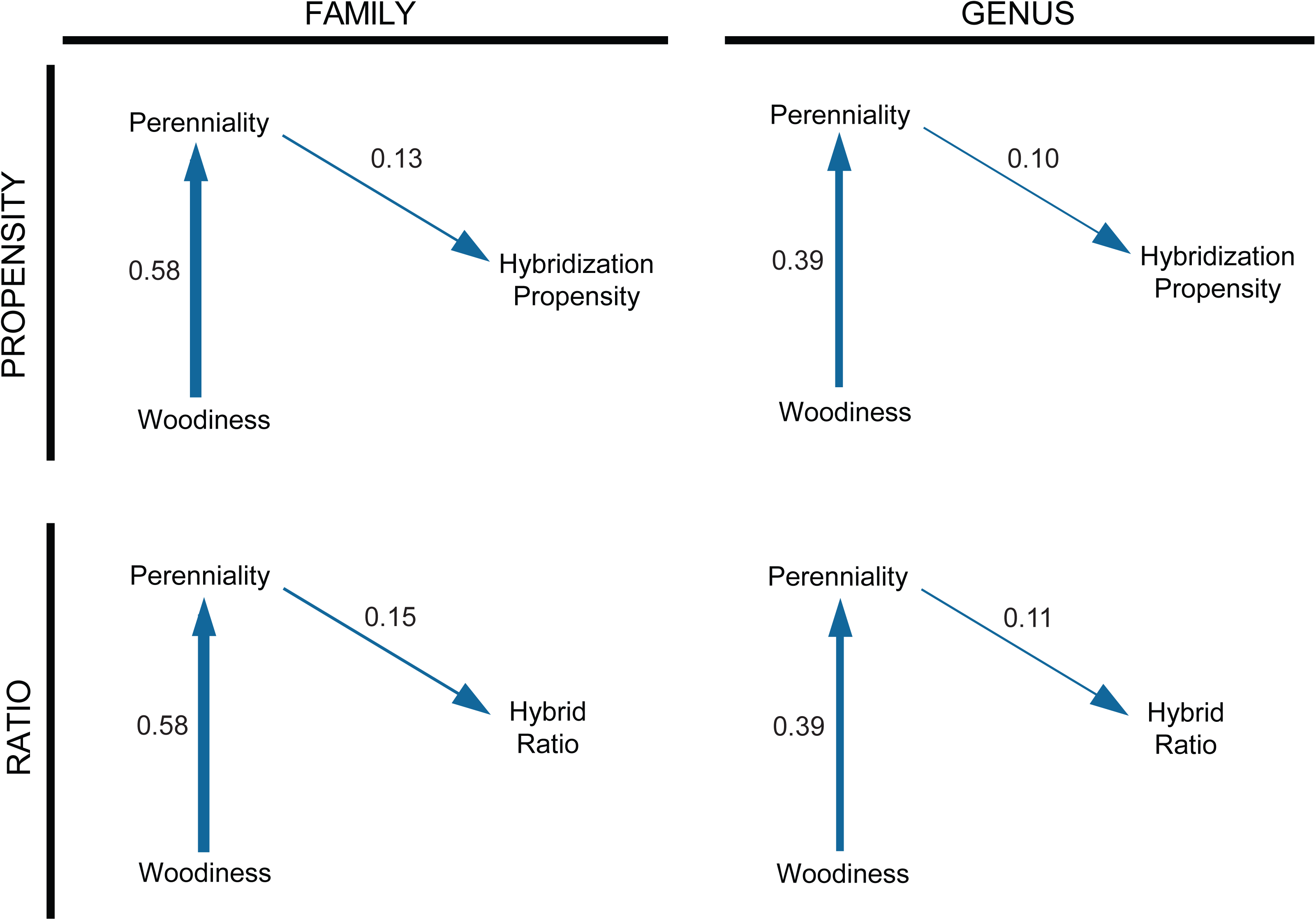
Predictors of hybridization propensity (top) and hybrid ratio (bottom) at the family (left) and genus (right) levels from phylogenetic path analysis using two predictors with large sample sizes that are also highly correlated: perenniality and woodiness. Final models were chosen via CICc from five candidate models (see Figure S1). Widths of the arrows indicate the strength of the coefficient and the direction of the relationship. Color indicates the sign (positive = blue, negative = pink) of the estimate. A lack of an arrow indicates that a relationship was not included in the best model.

### Raw correlations

For comparative purposes, we present raw correlations in a supplementary table. Several relationships between traits and hybridization rate or propensity were detected in the raw analyses that were not detected in the phylogenetically corrected analyses, emphasizing the importance of examining these relationships in a phylogenetic context (Table S4).

## DISCUSSION

Hybridization is not evenly distributed across the phylogenetic tree of life (Ellstrand *et al*. 1996), nor is it evenly distributed within plants, as we have documented here and elsewhere (Whitney *et al*. 2010). We detected several associations between hybridization rates and plant traits (perenniality, woodiness, outcrossing rate, pollination syndrome, reproductive system, genome size, and genome size variation) across the globe. Below, we organize our discussion of these associations sequentially, first discussing traits that may allow the *formation* of hybrids, followed by traits that may allow for the *persistence* of hybrids.

### Correlates of hybridization: factors that may allow for hybrid formation

Lineages may have detectable associations with specific factors that allow for the more frequent *formation* of hybrids. These associations may be direct or indirect in nature. For example, there may be a direct association between outcrossing and high levels of hybridization. High levels of outcrossing (or obligate outcrossing, as an extreme) mean that plants need to reproduce with another individual, necessitating the transfer of pollen, and increasing the odds of contacting and reproducing with another species when compared to selfing (Stace 1975, Ellstrand *et al*. 1996). Supporting this idea, we detected a trend for a positive association at the family level between outcrossing rate and hybridization propensity (Fig. 3, Table 3).

Other factors may be indirectly associated with hybridization. Grant (1958) hypothesized that associations between perenniality/woodiness and increased hybridization rates were actually indirect associations via outcrossing. He observed that perennial outcrossers were the most likely category of plants to participate in interspecific breeding and that autogamous or selfing plants were the least likely. We found associations between hybridization metrics and both woodiness and perenniality (Fig. 3, Table 3), and these traits were also correlated with outcrossing (Table S3). Our findings match previous hypotheses and non-phylogenetically corrected associations between hybridization and woodiness and/or perenniality (Stebbins 1959, Beddows and Rose 2018; Stace 1975, Ellstrand *et al*. 1996). In our analyses, the links between perenniality/woodiness and our hybridization measures were stronger than links with outcrossing rate (which had only a moderate association with hybridization propensity across families), but this discrepancy may be due to the restricted number of taxa for which we had outcrossing rate data (outcrossing data for 76 families and 158 genera, compared with perenniality and woodiness data for 195 families and 1754 and1767 genera, respectively, Table 2, Table S2). Perenniality and woodiness are positively correlated in plants, our evidence suggests that perenniality may be driving the association with hybridization, as there was more evidence for models including a direct path from perenniality to hybridization than a direct path from woodiness to hybridization (Fig. 4, Table S5, Table S6).

Factors not associated with outcrossing directly may also increase the chances of mating with heterospecifics and forming hybrids. Abiotic pollination syndromes may reduce pre-zygotic barriers to reproduction by allowing for promiscuous transfer of pollen, independent of biotic vectors. We found associations between abiotic pollination and hybridization at the family-level, but not the genus-level (Fig. 3, Table 3). We believe this is the first empirical dataset used to explicitly test for this association while correcting for phylogenetic non-independence (see Ellstrand *et al*. 1996, Rieseberg and Wendel 1993 for raw correlations, in both directions), and our results suggest that perhaps the less-discriminant abiotic pollination mode may lead to more hybridization. Additionally, low variation in genome size within a taxonomic group (which may signal the absence of ploidy variation) may be associated with the formation of hybrids, because ploidy barriers may block hybridization.

Interestingly, we failed to detect associations between hybridization and several hypothesized drivers. We (and others, Table 1, Table S1) posited that many of these traits would enable increased formation of hybrids via opportunity in sheer numbers or wide distributions (agricultural status, Red List status), or via reduced pre-zygotic barriers to hybrid formation (floral symmetry, breeding system). We note that the lack of detected associations could either be biologically real, or due to small sample sizes for some traits (Table S2). Further, other potential correlates not tested in this study could also promote the formation of hybrids (e.g., disturbance, low genetic divergence, Table 1).

### Correlates of hybridization: factors that may allow for hybrid persistence

Lineages may also have detectable associations with specific factors that allow for the *persistence* of hybrids once they have been formed. Early-generation hybrids are generally thought to exhibit either decreased fitness (hybrid breakdown) or, conversely, increased fitness (heterosis). The persistence of a hybrid lineage could be linked to either overcoming the latter or sustaining the former (stabilized heterosis). Long lifespans (associated with our traits perenniality and woodiness) may allow hybrid individuals with partial sterility to still have high levels of lifetime fitness, as a small number of viable seeds produced over multiple seasons can result in many offspring over time (Ellstrand *et al*. 1996). Thus, the association we detected between perenniality/woodiness and hybridization rate could be driven by effects on both hybrid formation (via outcrossing, see above) and persistence.

At the other extreme, heterosis due to heterozygosity at loci throughout the genome is expected to decline as sexual recombination results in the pairing of homozygous alleles in offspring (Conner and Hartl 2004). Stabilized heterosis is the preservation of the increase in fitness through time. Stabilized heterosis can be achieved through vegetative propagation, where early-generation fitness is maintained via the production of new individuals with a genetic composition identical to that of the parent. Consistent with this idea, we found that genera with more hybrids tended to have more vegetative reproductive systems (*vs.* generative) (Fig. 3, Table 3). There are several examples of clonal hybrids, for instance in *Tamarix* (Gaskin and Schaal 2002), *Myriophyllum* (Moody and Les 2002), and in many crop plants (reviewed in McKey *et al*. 2010).

Not all reproduction without outcrossing, however, is capable of preserving stabilized heterosis. For example, selfing (autogamy) should result in acceleration of the loss of heterosis due to a rapid reduction in heterozygosity (e.g., Johansen-Morris and Latta 2006). If a hybrid forms and then reproduces by selfing rather than outcrossing, it will not have the benefit of stabilized heterosis and the hybrid lineage may fail to persist. We found higher outcrossing rates in plant groups with more hybrids, perhaps reflecting this lack of hybrid persistence in selfing groups.

Some previous work in the genus *Cirsium* suggests that species with smaller genome sizes are more likely to form hybrids (Bureš *et al*. 2004). Although only marginally statistically significant, our evidence suggests a trend that groups with larger genomes can be associated with higher levels of hybridization propensity, contrary to this previous work. The association between larger genome sizes and higher hybridization rates could be due to the presence of numerous allopolyploids (hybrids produced from complete genomes of different species) within the group. Allopolyploidy could contribute to both high estimates of hybridization rates and large genome sizes for a given plant group, resulting in the observed associations. Further study is needed to investigate this pattern.

### Effects of taxonomic scale

Lineages that are more distantly related (longer time since divergence) tend to have stronger reproductive barriers between them than lineages that are more closely related (less time since divergence) (Coyne and Orr 1989 1997; Moyle and Nakazato 2010), although there are exceptions and this pattern may be dependent on other aspects of taxonomic scale (Moyle *et al*. 2004; Scopece *et al*. 2008; Nosrati *et al*. 2011). The majority of plant hybridization takes places within genera (Whitney *et al*. 2010), although instances of intergeneric hybridization have been observed, especially in non-flowering plants (Wagner *et al*. 1992; Wagner 1993; Fraser-Jenkins 1997; Garland and Moore 2012; Arrigo *et al*. 2013; Larson *et al*. 2014; Rothfels *et al*. 2015). We collected data at the generic level and analyzed these data at both the family (weighted) and genus taxonomic levels. Regressions tended to be more well-supported at the generic level after accounting for multiple comparisons (Fig. 3, Table 3). We found no well-supported relationship at one taxonomic level that was well-supported in the opposite direction at the other taxonomic level. Relationships found at the generic level and not found at the family level (for instance, between hybridization rate and reproductive system) could be due either to sample size differences (a statistical explanation) or the facts that genera within families differ with respect to specific traits, and that hybridization largely takes place within genera (a biological explanation). Relationships supported at the family level and not found at the genus level (for instance, between hybridization and pollination syndrome) could be due to increased precision in estimating both trait values and hybridization metrics within families, as the latter contain greater numbers of species than do genera.

### Measures of hybridization

Our measures of hybridization were based on the number of unique hybrid combinations produced, either as a proportion of potential hybrid combinations or simply using the number of nonhybrid species as a denominator. Our findings using both hybridization propensity and hybrid ratio were largely consistent. Not only were they significantly correlated at both the family and genus levels (Table S4) but their relationships with our proposed plant attributes were largely consistent. There were differences in significance when examining one or the other, but the trends were similar (Fig. 3, Table 3). We note that there is another metric which we did not employ, hybridization frequency, which takes into account the fraction of hybridizing parental species rather than their resultant taxa (Mallet 2005, Beddows and Rose 2018). Our database was constructed following Ellstrand *et al*. (1996) in a way that does not allow for the implementation of this metric, as we did not keep track of parental species. However, we note that the three hybridization metrics can be highly correlated (e.g., Beddows and Rose 2018) and thus suggest that analyses using hybridization frequency may not detect patterns different from those we report.

### Limitations

Although this study examines published floras that span three different continents and one island group, our conclusions may be limited and biased by the geographic extent examined. All but two of our floras are from Europe and mainland North America, with the Victoria, Australia and Hawai’i floras representing the Pacific Region. Four of the floras are from mainland North America, and these include almost half of all species observations (Table S1). In order to expand this dataset to other regions, we need comprehensive regional floras that specifically record instances of hybridization. Such floras are difficult to find, as they require both interest in hybrids by the authors and the decision to include information on them in the floristic treatment.

We collected data on hybridization using a method suited to their detection in regional floras. There is increasing evidence for instances of hybridization that are not necessarily morphologically apparent but are inferred using genetic or molecular evidence (i.e.: Cronn and Wendel 2004; Soltis *et al*. 2007; McIntosh *et al*. 2014; Mitchell and Holsinger 2018). At present, a comprehensive analysis including cryptic hybrids is not feasible, but as molecular methods become increasingly common (reviewed in Taylor and Larson 2019), a re-analysis incorporating expanded means of detecting hybrids would surely provide further insights.

## Conclusions

We found several strong phylogenetically informed associations between hybridization rates and plant attributes. Perenniality and woodiness across taxonomic levels, higher outcrossing rates and abiotic pollination syndromes at the family level, and less variable genome sizes at the genus level all associated with increased hybridization metrics and may be acting by increasing the formation of hybrids. Additionally, the associations between increased hybridization and perenniality, woodiness, outcrossing, and genome size, as well as more vegetative reproductive systems at the genus level, may be due to these factors increasing the persistence of hybrids that have already formed. We recognize that this evidence is correlational in nature and does not provide any causal inferences. Moreover, the explanatory power of our models was low (as measured by adjusted R^2^ values, Table 3). We caution that while we detected significant statistical associations, the vast majority of variation in hybridization rates remains unexplained. Future work is needed to experimentally test the nature of the relationships that we present here on a global scale. For instance, experiments comparing the evolutionary trajectories and population dynamics of closely related species pairs that are either abiotically or biotically pollinated (or both, such as ambophilous plants) could detect differences in rates of hybrid formation, and thus could support our correlative data. Our findings provide strong hypotheses for further investigating the drivers of hybridization and will aid in not only understanding hybridization as a stand-alone phenomenon, but also its role in invasion, range expansion, speciation, radiation, and diversification.

## ACKNOWLEDGEMENTS

Funding for the study has been provided by NSF DEB 1257965 and UNM startup funds (both to K.D.W.). Thanks to Loren Albert for assistance with trait scoring and to the Whitney-Rudgers lab members and four anonymous reviewers for feedback. This study has been supported by the TRY initiative on plant traits (http://www.try--db.org). The TRY initiative and database is hosted, developed, and maintained by J. Kattge and G. Bönisch (Max Planck Institute for Biogeochemistry, Jena, Germany). TRY is currently supported by the DIVERSITAS/Future Earth and the German Centre for Integrative Biodiversity Research (iDiv) Halle-Jena-Leipzig.

## AUTHOR CONTRIBUTIONS

K.D.W. and L.G.C. conceived of the original study. K.D.W., L.G.C., N.M., J.R.A., and K.C.P. collected data, and A.B.G. contributed data through the TRY database. N.M. performed the analyses. N.M. and K.D.W. wrote the manuscript. All authors contributed to revisions.

## DATA ACCESSBILITY

All hybridization data and phylogenetic trees are available from the Open Science Framework digital repository: doi:XXX.

