## Supplementary figures and images for "Correlates of hybridization in plants"

### Fig. S1

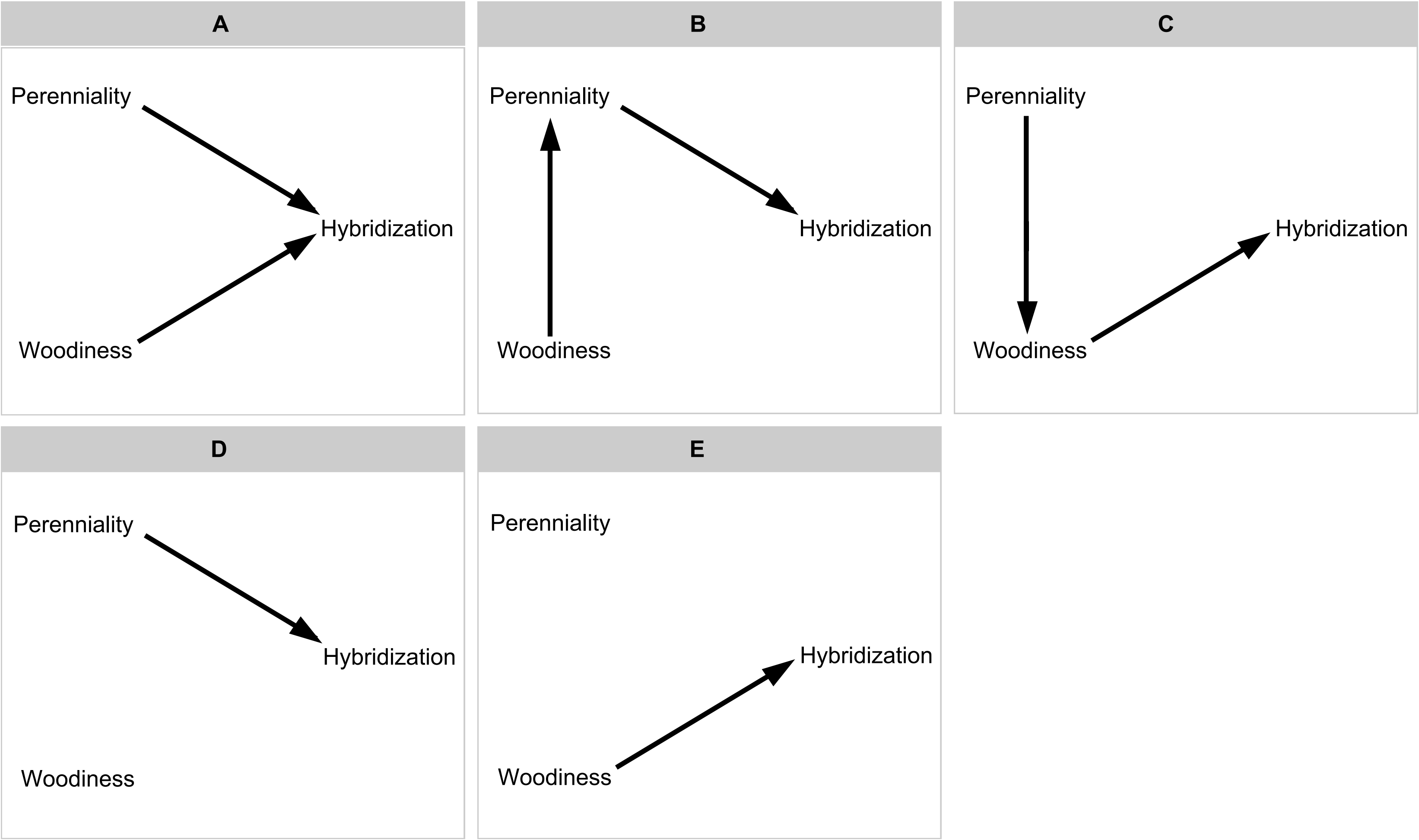
